# Climate trends and behavior of an avian forest specialist in central Amazonia indicate thermal stress during the dry season

**DOI:** 10.1101/2021.04.29.442017

**Authors:** Vitek Jirinec, Cameron L. Rutt, Elisa C. Elizondo, Patricia F. Rodrigues, Philip C Stouffer

## Abstract

Rainforest loss threatens terrestrial insectivorous birds throughout the world’s tropics. Recent evidence shows these birds to be declining in undisturbed Amazonian rainforest, possibly due to climate change. Here, we first addressed whether Amazonian terrestrial insectivores were exposed to climate change using 38 years of climate data. We found that climate has changed in central Amazonia, especially in the dry season, which was ∼1.3°C hotter and 21% drier in 2019 than in 1981. Second, to test whether birds actively avoided hot and dry conditions, we used field sensors to identify ambient extremes and prospective microclimate refugia within undisturbed rainforest from 2017 – 2019. Simultaneously, we examined how tagged Black-faced Anthrushes (*Formicarius analis*) used this space. We collected 1.4 million field measurements quantifying ambient conditions in the forest understory, including along elevation gradients. For 11 birds, we obtained GPS locations to test whether birds adjusted their shelter use (*n* = 2,724) or elevation (*n* = 640) across seasonal and daily cycles. For four additional birds, we collected >180,000 light and temperature readings to assess exposure. Field measurements in the modern landscape revealed that temperature was higher in the dry season and highest on plateaus. Thus, low-lying areas were relatively buffered, providing microclimate refugia during hot afternoons in the dry season. At those times, birds entered shelters and shifted downslope, reducing their thermal exposure by 50%. Because climate change intensifies the hot, dry conditions that antthrushes avoid, our results are consistent with the hypothesis that climate change lowers habitat quality for terrestrial insectivores. This sensitivity may be related to their declines within ‘undisturbed’ Amazonian rainforest.

## INTRODUCTION

Amazonia is the largest tropical forest and harbors a substantial portion of global biodiversity. At over 7 million km^2^, nearly equivalent to the continent of Australia, it contains ∼10% of described vertebrate species (IUCN, 2020; Silva et al., 2005). Rampant deforestation in the region has motivated research on the consequences of clearing and fragmentation for rainforest biota, including the avifauna (Bierregaard & Gascon, 2001; Peres et al., 2010; Stouffer, 2020). Ground-foraging insectivores are consistently among the species that respond most strongly to landscape disturbance. For example, experimental forest isolation led to attrition of terrestrial insectivores from fragments in central Brazil, with extinctions inversely proportional to fragment size (Stouffer & Bierregaard, 1995; Stratford & Stouffer, 1999). Similar patterns materialized in Ecuador (Canaday, 1996; Canaday & Rivadeneyra, 2001) and elsewhere in the Neotropics (Sekercioglu et al., 2002; Sigel et al., 2006, 2010; Visco et al., 2015). Strong sensitivity to forest disturbance thus makes terrestrial insectivores representatives of biodiversity within intact rainforest.

New studies have uncovered declines of terrestrial insectivores in what should be undisturbed Amazonia. In Ecuador, abundance of these species decreased markedly over a 14-year interval (Blake & Loiselle, 2015). Almost 2000 km away, a similar trend was recently described from the central Amazon: over four decades, terrestrial insectivores vanished from over half of primary forest sites and their relative abundance dropped substantially (Stouffer et al., 2020). In that study, terrestrial insectivores declined fastest among 12 ecological guilds examined, followed closely by near-ground insectivores. To document such population trends is very challenging; it requires long-term sampling using standardized methods to survey elusive species (Robinson et al., 2018). Within Amazonia, these two studies comprise the best available information on population trends of rainforest birds in the absence of direct disturbance. Alarmingly, both suggest that ground-foraging insectivores—already sensitive to landscape processes—are in trouble.

Why are terrestrial insectivores disappearing from intact forest? Research in disturbed landscapes offers a place to start. Hypothesized explanations range from vulnerability to changing forest structure (Laurance et al., 2002; Stratford & Stouffer, 2015), reduction in forest patch area (Stouffer, 2007), and several other factors (Powell et al., 2015; Visco et al., 2015). Notably, the ‘microclimate hypothesis’ posits that non-forest areas and forest edges harbor altered microclimates (*sensu* Chen et al., 1999) that are unsuitable for terrestrial insectivores, which are associated with shaded, cool, and wet conditions within forest interior. Isolated forest patches gain abnormal microclimate as a consequence of edge effects—they become brighter, hotter, drier, and these conditions grow more variable (Laurance et al., 2002). Anomalies were identified as most detrimental to organisms in stable environments, because ecological theory predicts that physiology is shaped by the conditions under which it evolved (Janzen, 1967). This notion is supported both for endotherms and ectotherms, as animals that display the narrowest physiological tolerances tend to reside in the tropics (Huey et al., 2009; Porter & Kearney, 2009), where temperature and precipitation are relatively stable throughout the year. Within Amazonia, terrestrial insectivores inhabit the most stable of environments—the floor of the forest interior, removed from both edge effects and the hotter, brighter, drier canopy >20 m above the ground. Here, temperature and light intensity near forest edge and the canopy climb, while the reverse occurs for water availability (Kapos, 1989; Scheffers et al., 2013; Sheldon et al., 2018; Walther, 2002). Thus, in the absence of edge effects, pristine forest appears to deliver optimal environment for terrestrial insectivores.

However, even large tracts of primary forest may be disturbed. Human activities cause the climate to diverge from historical norms across the globe. In the tropics, climate models forecast abnormally high temperatures (Mora et al., 2013; Neelin et al., 2006) and more temperature fluctuations (Bathiany et al., 2018). In Amazonia, average temperature has climbed ∼0.05°C year^-1^ since 1973 (Almeida et al., 2017). In contrast with consistent warming, precipitation is more spatially variable. Shifts in rainfall regimes are sometimes manifested through wetter wet seasons, but dry seasons are often drier and longer (Almeida et al., 2017; Fu et al., 2013), with droughts predicted in the future (Neelin et al., 2006). Climate change has already been linked to changes to forest structure by amplifying tree mortality, abundance of dry-affiliated species, and biomass of lianas (Aleixo et al., 2019; Esquivel-Muelbert et al., 2019; Laurance et al., 2014). Evidence is hence mounting that today’s terrestrial insectivores occupy a hotter and drier Amazonia than just a few decades ago, with conditions increasingly aligned with forest fragments.

The Biological Dynamics of Forest Fragments Project (BDFFP) offers the opportunity to assess the impacts of climate change on terrestrial insectivores. Located just north of Manaus, Brazil (Figure 1), the BDFFP is the nexus of research on the Amazon rainforest (Laurance et al., 2018), including birds (Stouffer, 2020), and is one of the sites where terrestrial insectivores declined in primary forest (Stouffer et al., 2020). Here, we address two objectives while drawing on extensive research on terrestrial insectivores in local disturbed and undisturbed landscape. First, we assess evidence for exposure of terrestrial insectivores to climate change *in situ* using 38 years of data from a climate reanalysis focused on our study site rather than broad regional patterns. Second, to test our hypothesis that hot and dry conditions are unsuitable for birds, we use field sensors to identify cyclic periods of ambient extremes and prospective microclimate refugia, while simultaneously tracking the behavior of Black-faced Antthrush (*Formicarius analis*), a terrestrial insectivore. These results reveal that birds emblematic of undisturbed Amazonia attempt to avoid the very conditions which climate change exacerbates within primary forest thought to be intact.

**Figure 1.**
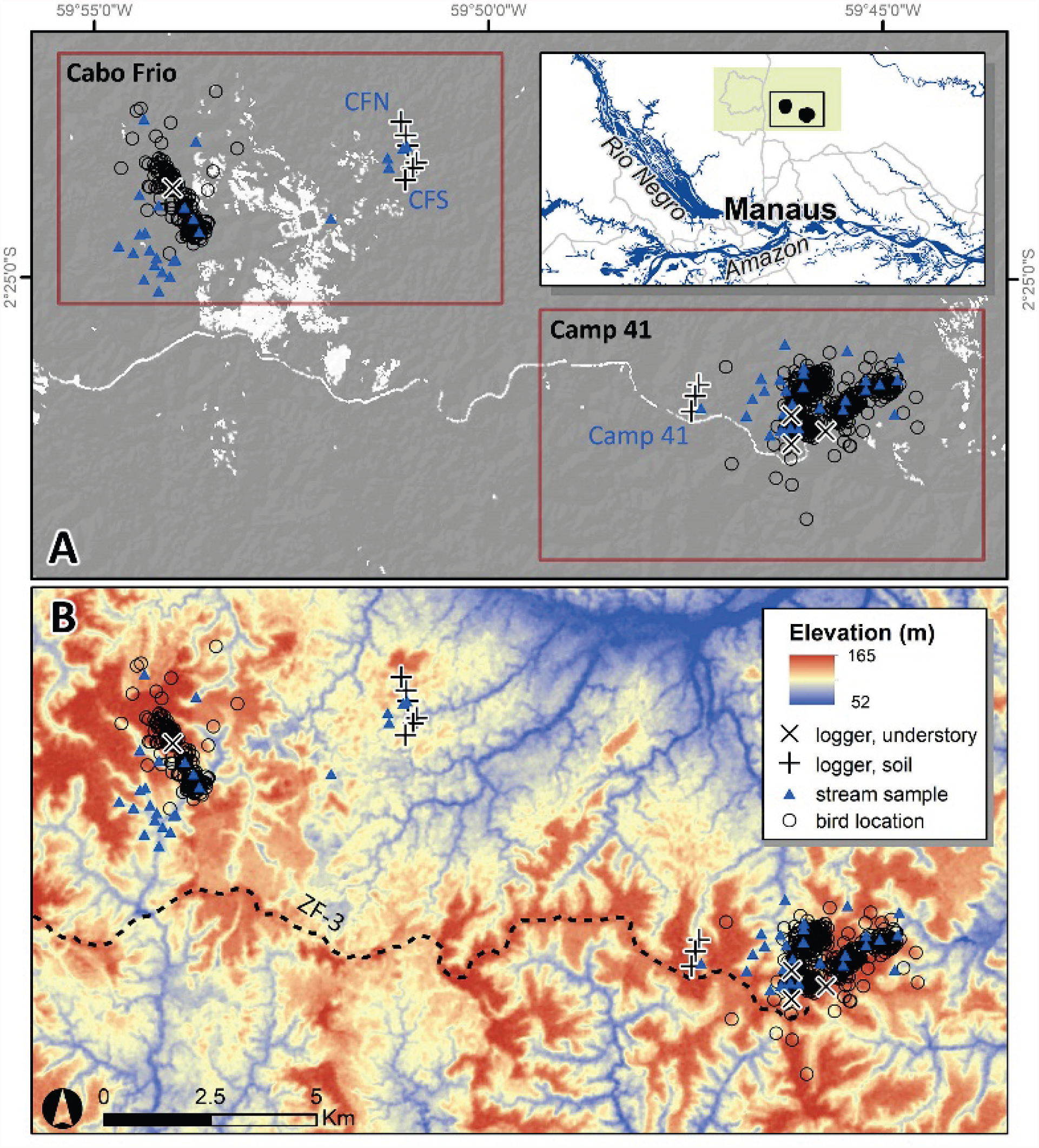
Survey areas at the Biological Dynamics of Forest Fragments Project in the state of Amazonas, Brazil. Panel ‘A’ depicts the two study areas, Cabo Frio and Camp 41, with Landsat-derived forest cover in 2017 (gray). Panel ‘B’ shows topographical variation (range ∼100 m) in the same scene. Both panels contain the 640 GPS locations of 11 *Formicarius analis* individuals considered in this analysis (2017 – 2019), as well as the locations of sensors that measured ambient conditions. The green rectangle (inset) shows the region where we summarized ERA5- Land data for the climate change analysis.

## MATERIALS AND METHODS

### Study area, seasonality, and climate

The BDFFP typifies forests of lowland Amazonia. The project spans approximately 1,200 km^2^ of which ∼95% in 2017 was covered with *terra firme* forest (Rutt et al., 2019), growing on nutrient-poor soils of the Guiana Shield (Gascon & Bierregaard, 2001). We collected data within continuous primary forest at the Cabo Frio and Camp 41 camps (Figure 1). Both areas have topography characterized by moderate relief where stream valleys dissect upper-elevation plateaus often located ∼100 m higher, creating a highly reticulated landscape (Figure 1B).

Temperature and precipitation are high throughout the year but vary seasonally. The region experiences an annual cycle of predictable rainfall fluctuation (Fu et al., 2013). The main goal of this study was to assess whether birds are sensitive to hot and dry conditions; we delineated the seasonal cycle at the BDFFP to understand when birds are expected to respond. To do so, we compiled climate data for 38 years from the ERA5-Land reanalysis (Copernicus Climate Change Service, 2017) for the area where birds were historically sampled (Stouffer et al., 2020). ERA5-Land uses satellite and land-based records with a climate model to estimate several climate values at 9 km resolution. We summarized precipitation and temperature at 2 m above ground level within ∼4,000 km^2^ surrounding the study sites (Figure 1), yielding monthly summaries from 1981 to 2019, and then used generalized additive models (GAMs; see *Statistical analyses*) to quantify the annual climate cycle. Our models revealed August to be the middle of the dry season, when precipitation is about a third of the total rainfall in March and April, the wettest months (Figure 2). Temperatures peak slightly after the driest period—they are ∼1.5°C higher around October than in the middle of the wet season (Figure 2).

**Figure 2.**
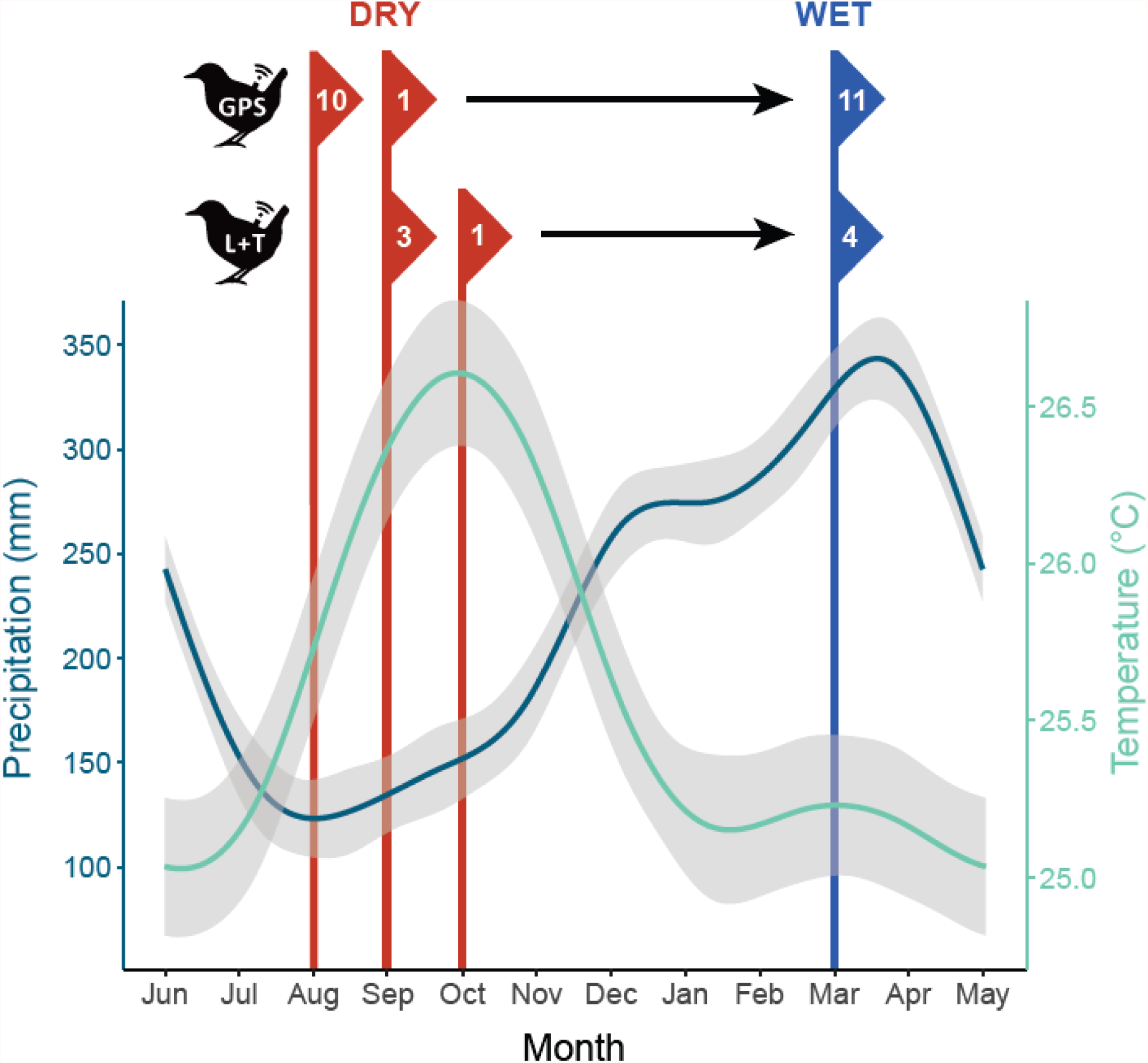
Annual climate and schedule of bird sampling. We first used an autoregressive generalized additive model and local rainfall and temperature data to broadly identify periods of annual climate extremes for comparison of bird behavior. Climate data come from ERA5-Land reanalysis and are a compilation of 38 years for our study area. Lines are model fits with the 95% CI shaded in gray. We then tracked Black-faced Antthrushes (*Formicarius analis*) over one annual cycle, either with a GPS (*n* = 11) or temperature logger (*n* = 4). For birds with GPS tags, individuals were tracked over one month in the dry season (Aug or Sep) and one month the following wet season (Mar), depending on year. Birds with biologgers were tracked over two months in each season, starting either in Sep, Oct, or Mar.

Is the BDFFP experiencing climate change? To answer this question, we grouped the ERA5-Land monthly summaries by season: Jun – Nov (dry) and Dec – May (wet). These seasonal classifications were based on our within-year analysis (Figure 2) and follow past designations for the BDFFP (Stouffer et al., 2020). Climate covariates were modeled with GAMs to determine whether climate change occurred at our site and if climate extremes varied by season.

### Microclimate refugia

Ambient extremes can manifest in time and space. Seasonal climate suggests that extremes generally occur during the dry season at the BDFFP (Figure 2). Ambient conditions thus often differ the most between about August and March—intervals that define the trough and crest of the annual rainfall and temperature cycles. Dry season, and especially dry season afternoons, are generally the hottest and driest periods in Amazonia. However, these extremes may be reduced in microclimate refugia—localized areas that remain relatively cool and wet. For example, in primary rainforest in the Philippines, the rate of temperature change in dense vegetation and tree cavities was 43% relative to ambient temperature, and exposure to extreme temperatures occurred over intervals that were 22 times longer (Scheffers et al., 2014). In addition to this physical shelter, topography may also buffer ambient extremes. Temperature drops with increasing elevation at large scales—a fact of little consequence in most of low-elevation Amazonia. In contrast, at the BDFFP landscape, valleys likely remain wetter and cooler. The bottoms of these ‘micro-catchments’ usually contain streams that are maintained by groundwater throughout the year (Epron et al., 2006; Tomasella et al., 2008). Aside from direct cooling, water buffers against temperature change (Davis et al., 2019; Elsenbeer, 2001; Fridley, 2009), and wetter soils allow evapotranspiration to occur when drier areas may lead to stomatal closure (Aleixo et al., 2019). Hence, we expected ambient extremes to arise in the afternoons of the dry season, with physical cover and valley bottoms offering buffered microclimates.

We empirically identified microclimate refugia with field measurements. The buffering property of physical cover in a rainforest setting was revealed elsewhere (Scheffers et al., 2014); here we focused on elevational refugia within Amazonian micro-catchments (Tomasella et al., 2008). To determine whether valleys buffered ambient extremes, we measured temperature and soil moisture along three elevational transects with a total of nine datalogging stations (Figure 1, Appendix 1). We selected transects along slopes in primary forest based on LiDAR elevation data, and placed stations away from treefall gaps such that each transect held one station at the valley bottom (level 1), mid-slope (level 2), and plateau (level 3). The nine stations were thus evenly classified by elevation, embedded within transects spanning ∼600 m of length and ∼40 m in elevation (Appendix 1). Each station contained one logger (TrueLog100) and one sensor (SMT100), both manufactured by Truebner (Truebner GmbH, Neustadt, Germany). We inserted sensors fully into the ground; temperature and water content readings thus correspond to the topmost 11 cm of soil—a relevant stratum for birds that seldom leave the forest floor. Loggers were programmed to measure temperature (°C) and soil moisture (% volumetric water content) every 10 min for the duration of sampling. We assigned the sampling periods to be Aug – Nov (dry season; DS) and Feb – May (wet season; WS). We chose these months because they fell within the DS or WS based on historical rainfall data (Figure 2), contain the most bird observations, and are of equal length for the analysis of microclimate conditions. Aside from automated measurements at these nine locations, we manually sampled stream temperature across a broader area. These 53 samples were collected evenly throughout daylight hours over 32 days within 21 Jun – 13 Sep 2019 in the two study areas (Figure 1) and represented the effect of stream temperature on valley-bottom microclimate in the DS.

### Tracking bird behavior

We expected birds to seek microclimate refugia during the DS, as vagile animals can exploit heterogeneity within their habitat to maintain optimal body temperature by behavioral thermoregulation (Cowles & Bogert, 1944; Huey et al., 2003; Porter et al., 1973; Stevenson, 1985). To assess whether this occurred, we used the WS as the baseline for comparison, and tested our prediction that birds used shelter more and shifted to lower elevations in the DS.

We selected *Formicarius analis* as a representative terrestrial insectivore. The BDFFP contains 13 species of terrestrial insectivores (Stouffer, 2007), but only *F. analis* is both adequately common and sufficiently large to carry GPS devices (Johnson & Wolfe, 2017; Rutt et al., 2017). The species is a permanent resident that maintains a year-round territory defended by a mated pair, but territory stability can fluctuate among years (Stouffer, 2007). As with nearly all terrestrial insectivores, capture rates of *F. analis* displayed a declining trend since the early 1980s (Stouffer et al., 2020).

We caught territorial birds with target-netting. First, we located birds by broadcast of conspecific playback to elicit a vocal response of a local territory owner. After detection, we set two or more 12-m mist nets (32 mm mesh) arranged in a ‘V’ formation with the playback in the center. When the focal bird approached playback, a concealed operator then flushed the bird into nets. We then repeated the capture process the following year to recover devices. Initial trapping and data recovery required 257 days (136 at Camp 41 and 121 at Cabo Frio) spread across 12 months in three years: 2017 (Jun – Aug), 2018 (Jun – Oct), and 2019 (Jun – Sep). At each capture occasion, we took standard morphometric measurements and outfitted birds with either GPS or biologging tags. Because age and sex is difficult to determine for this species (Johnson & Wolfe, 2017), we assumed all captured birds were adult males due to their territorial responses to song playback. For details about tag fitting, materials, and the lack of detrimental effects of tagging on study birds, see Jirinec et al. (In press).

We tracked bird positions in space and time with archival GPS tags (PinPoint-50; Lotek, Newmarket, Ontario, Canada). Tag configuration (‘SWIFT’ fixes) maximized the number of locations at the expense of greater location error and occasionally large outliers. SWIFT fixes involved a 12-second timeout after which devices either acquired sufficient GPS signal to determine location (fix = success) or not (fix = fail). We programmed tags to attempt four fixes a day (07:00, 10:00, 13:00, 16:00 local time) over one month in each of the two seasons. This scenario allowed us to compare a similar number of locations (up to 124 fixes per season) for each bird. In 2017 and 2018, we deployed a total of 18 GPS tags, of which we recovered 11 in subsequent years. Recovered tags contained locations from eight birds in the 2017 DS – 2018 WS cycle and three birds in the 2018 DS – 2019 WS cycle (Figure 2). Using successful fixes, we extracted elevation from a 12-m WorldDEM (Riegler et al., 2015) to determine whether birds reduced their exposure to ambient extremes by moving downslope.

While we used successful GPS fixes to measure elevation shifts, we applied both successful and failed fixes to track shelter use. Because the precision of GPS tags was insufficient to resolve microhabitat directly from x-y coordinates, we employed the inverse probability of GPS fix as a proxy for shelter use. Dense vegetation and other physical barriers hinder satellite signal, leading to relatively fewer locations within these areas (Di Orio et al., 2003; Jiang et al., 2008; Recio et al., 2011). Although this reality is usually undesirable, here it allowed us to track shelter use indirectly when we standardized the intervals over which tags search for satellites (12 seconds). Because GPS signal varies little spatially and temporally (*GPS.Gov: Space Segment*, 2020), the probability of successful GPS fix is inversely proportional to the level of surrounding obstruction, with probability of fix near ‘1’ in open sky and ‘0’ inside logs, stumps, or dense vegetation. To help validate this assumption, we radio-tracked a single *F. analis* with VHF telemetry between 19 Jul and 24 Aug 2017, which allowed us to observe behavior directly. If birds sought cover during ambient extremes, those times should coincide with the lowest probability of GPS fix.

To quantify exposure to ambient conditions, we tracked birds with biologging tags (‘geolocators’; Intigeo-P65B1-11T-20deg, Migrate Technology, Cambridge, UK). Tags recorded light intensity (lux) every 5 min and temperature (°C) every 15 min for two months each season. Biologgers sampled light intensity atop a stalk to prevent feather shading, while thermometers were embedded in tags that were flush with the bird’s body (Appendix 2). Thus, light readings represented direct exposure to light, while temperature was a combination of body and ambient temperature. To better understand the relationship between tag measurements and ambient conditions, we sampled light and temperature with four biologgers that were placed near tagged birds ∼10 cm high in the understory of mature forest, at a mean elevation of 136 m (Figure 1, Appendix 2). We also measured the body temperature of 36 unique birds by cloacal measurements with a medical thermometer (upper temperature limit = 43.0°C). Measurements were taken as soon as possible after capture to lessen any possible effect on body temperature (Prinzinger et al., 1991). In 2017 and 2018, we deployed 13 biologgers on birds, of which we recovered four in the following years. Light and temperature data came from three birds tracked over the 2017 – 2018 seasonal cycle: Sep – Oct and Mar – Apr, and one bird tracked over the 2018 – 2019 cycle in Oct – Nov and Mar – Apr (Figure 2). Ambient loggers collected data in concert with bird tags, providing an approximation of ambient conditions that birds were exposed to. This allowed us to test for behavioral thermoregulation by comparing measurements from identical devices on birds and in the environment.

### Statistical analyses

To analyze climate and biologger data, we employed generalized additive models (GAMs) implemented in R package ‘mgcv’ (Wood, 2020). GAMs are similar to linear models, but allow for modeling of non-linear relationships in the response using smooth functions of explanatory variables (Wood, 2017). Package ‘mgcv’ calculates an approximate p-value for the smooth term; a low p-value constitutes evidence for a significant relationship—either positive or negative—somewhere along the range of the covariate, and plot of the smooth should be examined to interpret results (Wood, 2017).

To quantify annual seasonality and to test for climate change with ERA5-Land at the BDFFP, we used GAMs that accounted for autocorrelated observations. We selected the cyclic cubic regression spline basis (bs = ‘cc’, k = 10) for annual seasonality models and the thin plate regression spline basis (bs = ‘tp’, k = 10) for climate change models; all were fit with restricted maximum likelihood and Gaussian distribution. We specified generalized additive mixed models with autoregressive moving average (ARMA) error structure for each climate variable, modeled as the smooth of the numeric month in annual seasonality models and as the smooth of year in climate change models. We set ARMA within each year and selected the AR order by first plotting residual autocorrelation functions of non-ARMA models, followed by iterative likelihood ratio tests among models with plausible AR orders to select the appropriate model structure.

Last, to analyze biologger data, we used different GAM structures for light and temperature with separate seasonal analyses. For light exposure, we modeled light intensity as the smooth of numeric time of day (0 – 23.99) using the cyclic cubic regression spline (bs = ‘cc’, k = 50), the smooth of the successive observation rank per group (bird or logger) with the Gaussian process basis (bs = ‘gp’, k = 50), and group as the random effect (bs = ‘re’, k = 50). Light exposure models were fit with restricted maximum likelihood and Gamma distribution with a log link function. Temperature model structure was identical, although we used Gaussian distribution in place of Gamma.

To test whether valleys buffered ambient extremes, we calculated seasonal profiles by elevation using locally estimated scatterplot smoothing (LOESS). Prior to smoothing, we averaged timeseries by elevation (e.g., mean of the three valley transects per 10-min interval)— this resolved data gaps and created a single timeseries group per elevation. Each LOESS line represents the estimate of R function loess(), with α = 0.5 on the group average. For temperature and soil moisture, these were averages of the raw measurements. For daily temperature summaries, we first extracted the minimum, maximum, and range in temperature per day in the group average, and then estimated the LOESS lines of the summary timeseries.

Prior to modeling of bird GPS data, we quantified tag location error and incorporated it into our analyses. Vegetation and topography reduce not only the probability of fix but also the accuracy of acquired locations (Jiang et al., 2008), which was of particular concern for birds with SWIFT-enabled tags that often used valleys in mature rainforest. We therefore placed two calibration sensors (identical to bird tags) to represent stationary birds—they were positioned ∼10 cm high in the understory of mature forest, near study birds, and were stratified by elevation (lower tag: 91 m, upper tag: 140 m; Appendix 2). Although calibration tags sampled a somewhat different time of year, we programmed them with the same daily fix schedule as study birds, from 17 Jun – 31 Aug 2019. Out of 608 fix attempts, this yielded 411 successful locations, which we then used in R package ‘ctmm’ (Fleming et al., 2020) to estimate location error. Because satellite-based screening methods such as HDOP can be misleading and eliminate accurate locations (Recio et al., 2011), we instead identified spatial outliers for our calibration tags based on relative distance to other locations. For each tag, we first we used the outlie() function in ‘ctmm’ to calculate the distance from the median longitude and latitude to each point, which highlighted one obvious outlier for the lower tag (2,200 km deviation from core cluster) and another for the upper tag (1.2 km). We removed both of these outliers before estimating location error by running the uere.fit() function in ‘ctmm’ for each remaining calibration dataset. After generating an error radius of 52 m (95% CI: 49 – 56 m) and 41 m (38 – 44 m) for the lower and upper tag, respectively, we conservatively estimated that each valid bird fix was on average 52 m away from the true location. This horizonal error is relatively small, given the typical home range size (Appendix 2). We similarly removed outliers from the location dataset for each of the 11 GPS tags recovered from birds. As with the calibration data, we used outlie() to calculate core deviation for each bird and removed all locations that were farther than 1.5-times the interquartile range in core deviation. We believe that this vetting process constituted an objective way to remove outliers and quantify spatial error from small GPS tags deployed in this challenging environment.

To model bird GPS data, we used linear mixed models (LMMs) and generalized linear mixed models (GLMMs) in R package ‘lme4’ (Bates et al., 2020). First, to determine whether GPS fix probability varied temporally, we used GLMMs to model fix attempt as a binary response (success = 1, fail = 0) with season (dry, wet) and time of fix (hour: 07:00, 10:00, 13:00, 16:00) as explanatory variables and bird as a random effect. We applied the binomial error distribution with the logit link function (logistic regression) and fit these models by maximum likelihood with adaptive Gauss-Hermite quadrature using 25 sample points. This dataset involved all 2,724 fix attempts for the 11 recovered GPS tags, including failed fix attempts and successful fixes that were later removed as outliers. Second, we used the final valid location dataset (filtered via ‘ctmm’) to test whether birds shifted elevations during ambient extremes. To attach elevation to GPS data, we averaged DEM values with ArcGIS (ArcMAP 10.7.1, ESRI, Redlands, California, USA) within 52 m of each valid fix to account for estimated horizontal error. We then used LMMs to model elevation as the response with season and time of fix as explanatory variables and bird as the random effect. To compare the relative importance of each covariate, we built the same four models for elevation and the probability of GPS fix and compared competing models using AIC_c_ (Burnham & Anderson, 2002). The models were (a) a null (intercept-only), (b) hour, (c) season, and (d) an hour*season interaction. All models were fit using maximum likelihood, but we refit top LMMs with restricted maximum likelihood for interpretation of coefficients (Zuur et al., 2009). Finally, for each top model, we examined model diagnostics in R package ‘DHARMa’ (Hartig, 2020) using 10,000 simulations, as well as the model fit with conditional coefficients of determination (r^2^_c_) calculated with r.squaredGLMM() in R package ‘MuMIn’ (Bartoń, 2020). Analyses were conducted using R version 3.6.3 (R Core Team, 2020).

## RESULTS

### Ambient conditions

Climate has changed at the BDFFP over the last four decades. Within- and among-year variation in both temperature and precipitation was considerable, but three of the four models revealed significant climate trends over time (Figure 3). DS temperature had the strongest positive relationship with year—in 2019, DS temperature was ∼1.3°C higher than in 1981. Temperature in the WS also rose—in 2019, WS temperature was ∼0.6°C higher than in 1981. DS rainfall declined over time, with 2019 precipitation totaling ∼34 mm (21%) less than in 1981. WS rainfall indicated an increasing, but insignificant trend. Hence, terrestrial insectivores at the BDFFP are currently exposed to significantly hotter and drier conditions than in the early 1980s—especially during the DS.

**Figure 3.**
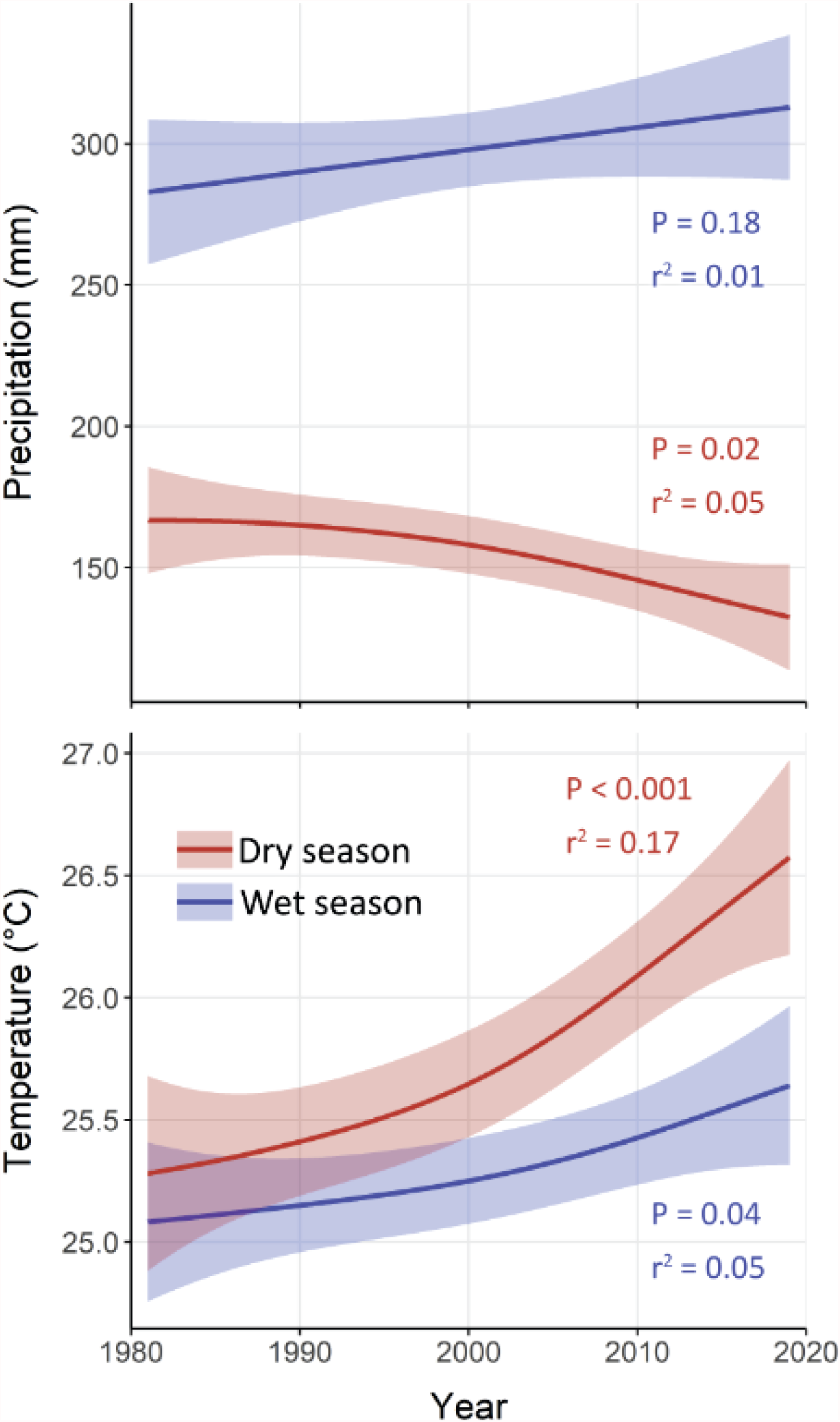
Climate change at the Biological Dynamics of Forest Fragments Project within Amazonian Brazil. Lines are fitted generalized additive models with ARMA errors, with shading representing 95% CI. Climate data are monthly summaries compiled for the BDFFP area from ERA5-Land reanalysis.

In modern forest, microclimate differed by time of day, season, and elevation (Table 1, Figure 4). The nine microclimate stations, operational Jun 2017 – Sep 2019, acquired 594,119 readings of both temperature and soil moisture over the four seasonal intervals. Only two gaps in the timeseries occurred, when rodents severed sensor cables, resulting in no data between 6 Nov 2018 – 21 Jun 2019 at CFN-3 and 11 Mar 2019 – 21 Jun 2019 at CFN-2. Although we originally placed all stations away from forest gaps, single trees fell within 10 m of both Camp 41-1 and CFS-1 during the WS of 2018. Soil moisture was lowest overall in the DS, but elevation created the strongest contrasts, with driest conditions at uppermost sites while valleys remained much wetter. Temperature varied by season, with higher mean, daily minimum, maximum, and range in DS. Extremes intensified at upper elevations. Daily peaks in temperature emerged most often between 14:00 and 15:00, regardless of season, although they tended to occur ∼30 min later in the DS and were more variable during the WS (Table 1). Stream temperature during the DS was relatively cool and varied little across space and time (mean ± SD = 24.6 ± 0.4°C, range = 23.6 – 25.7, *n* = 53). In short, afternoons in the dry season produced the hottest and driest periods of the year, but valleys offered microclimate refugia where conditions were milder.

**Figure 4.**
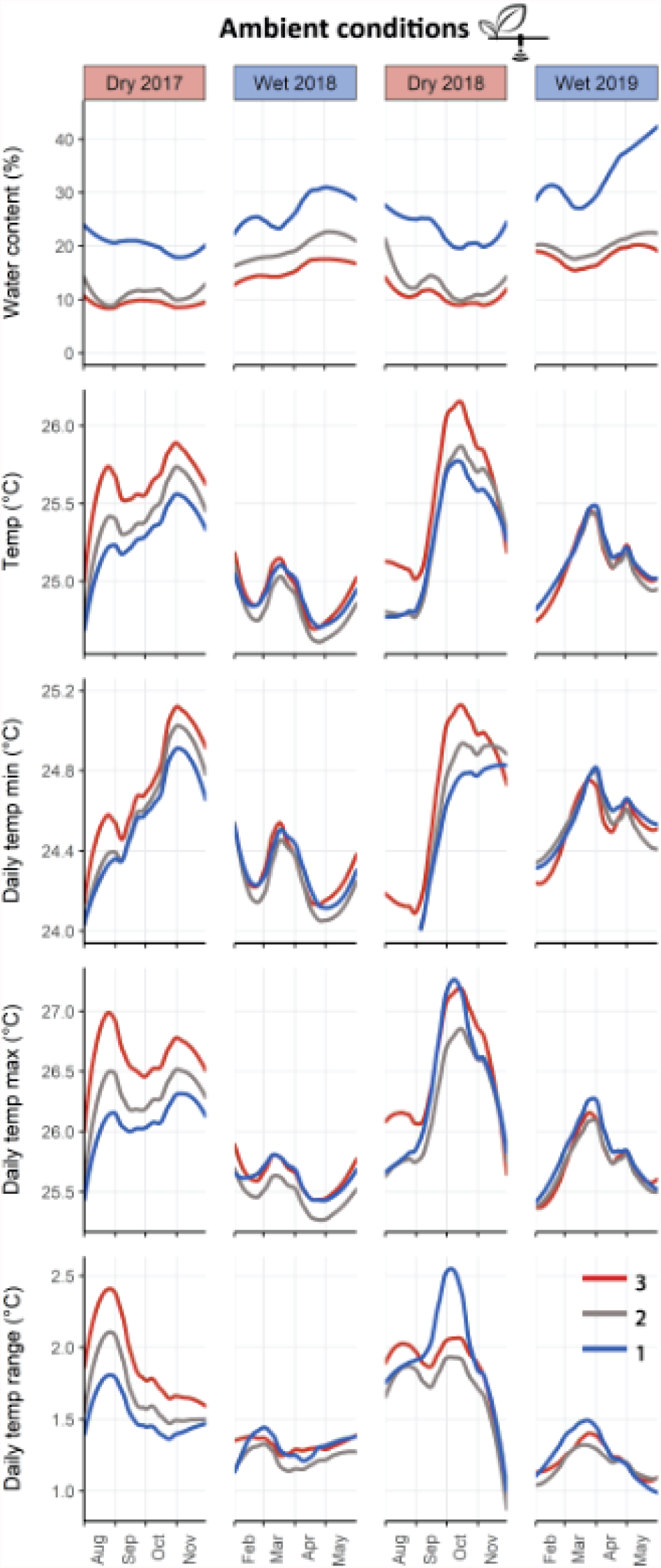
Forest floor microclimate within Amazonian micro-catchments in primary *terra firme* forest. Plots depict water content and temperature measurements from the top 11 cm of soil at nine microclimate stations arranged along three elevational transects, each with a station placed in a valley (1, blue), midway up a hillside (2, gray), and atop a plateau (3, red). Stations obtained a reading every 10 minutes during each four-month season; here we here show the locally estimated scatterplot smoothing of the average for each elevational group (i.e. each line smooths the mean of the three timeseries).

**Table 1.**
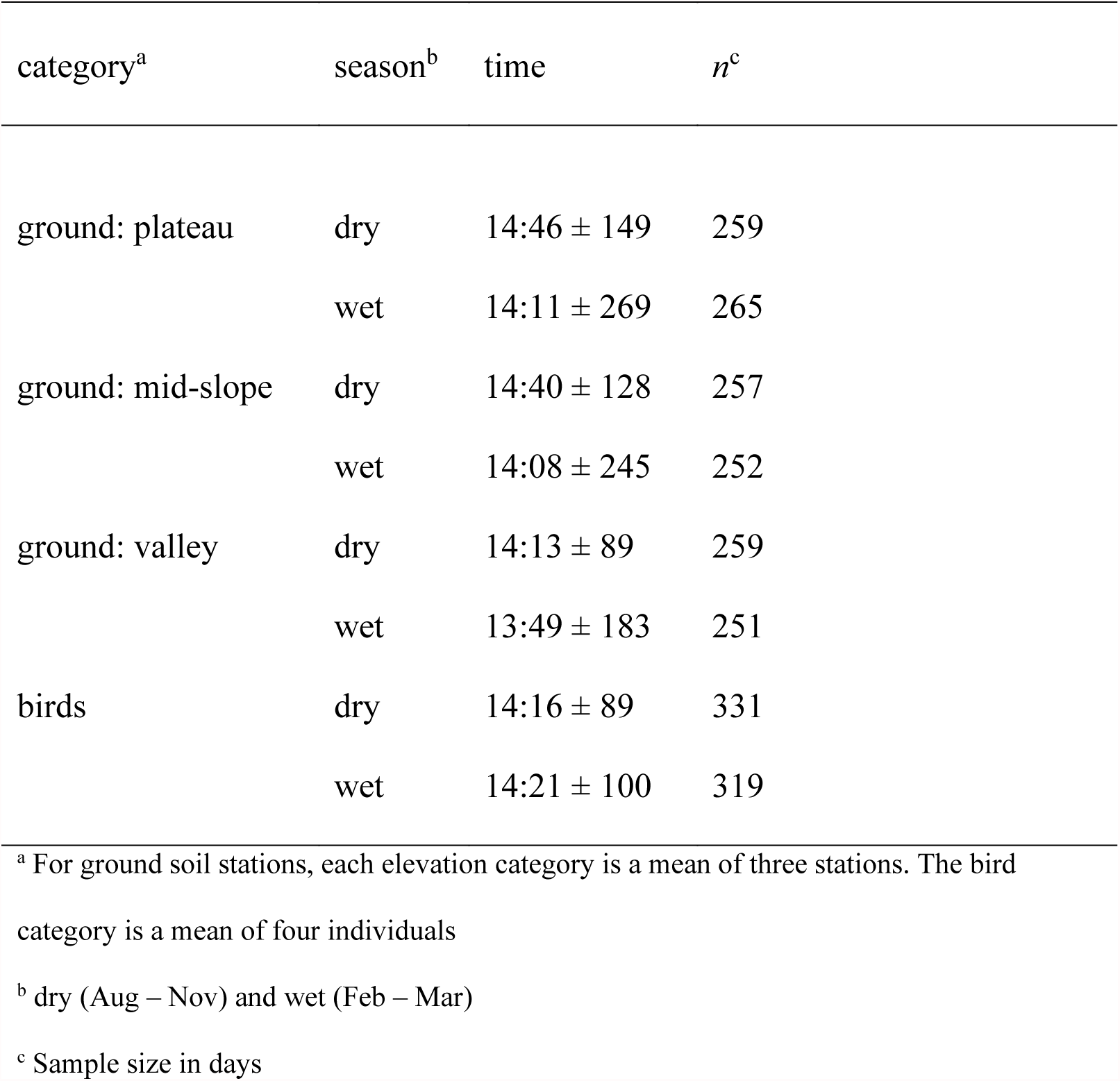
Timing of daily temperature extremes (mean ± SD in min) by season for ground microclimate stations and bird biologgers.

| category<sup>a</sup> | season<sup>b</sup> | time | *n*<sup>c</sup> |
| --- | --- | --- | --- |
| ground: plateau | dry | 14:46 ± 149 | 259 |
| | wet | 14:11 ± 269 | 265 |
| ground: mid-slope | dry | 14:40 ± 128 | 257 |
| | wet | 14:08 ± 245 | 252 |
| ground: valley | dry | 14:13 ± 89 | 259 |
| | wet | 13:49 ± 183 | 251 |
| birds | dry | 14:16 ± 89 | 331 |
| | wet | 14:21 ± 100 | 319 |

<sup>a</sup> For ground soil stations, each elevation category is a mean of three stations. The bird category is a mean of four individuals
<sup>b</sup> dry (Aug – Nov) and wet (Feb – Mar)
<sup>c</sup> Sample size in days

### Bird behavior

The 11 GPS tags attempted a total of 2,724 locations, with 688 (25%) successful fixes (640 [23%] after outlier removal). The four bird biologgers returned a total of 140,245 light and 46,748 temperature readings, whereas the four ambient loggers returned 125,232 and 41,744 readings, respectively.

We detected significant temporal patterns in the probability of GPS fix—our proxy for shelter use. The top model, by ΔAIC_c_ > 53, was the interaction model that contained both hour and season (Table 2). Using 07:00 in the WS as the baseline, the probability of GPS fix (inverse of shelter use) was significantly higher at 13:00 and 16:00 in the WS but dropped substantially at 13:00 and 16:00 in the DS (Figure 5A). In the seasonal model, probability of a successful fix was 27% in the WS and 22% in the DS (*β*_dry_ = -0.29, SE = 0.09, *P* = 0.001).

**Figure 5.**
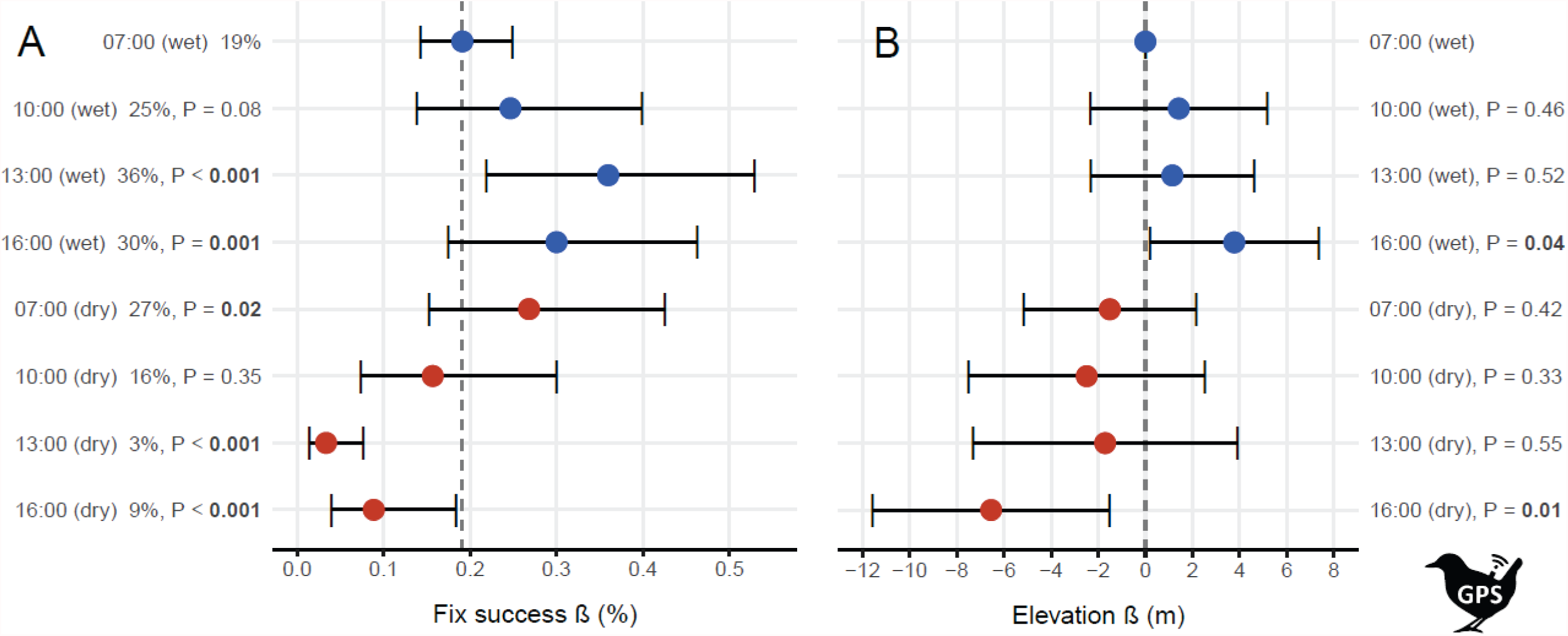
Probability of GPS fix (A) and elevation (B) by time of day and season for tags on 11 *Formicarius analis* in primary, continuous *terra firme* forest. Each bird was tracked four times daily over one month in the dry season (red) and one month in the wet season (blue) in 2017 – 2019. Error bars represent 95% confidence intervals and P-values test the difference against a baseline (dashed line: 07:00 in the wet season).

**Table 2.** Model selection for probability of GPS fix, including the number of model parameters (K), AIC_c_ weight (*w*), and log likelihood (LL).

| model | covariates | K | AIC<sub>c</sub> | ΔAIC<sub>c</sub> | *w* | LL |
| --- | --- | --- | --- | --- | --- | --- |
| interaction | hour*season | 9 | 2988.1 | 0 | 1.00 | -1485.0 |
| season | season | 3 | 3041.9 | 53.8 | 0.00 | -1518.0 |
| diel and season | hour + season | 6 | 3044.6 | 56.4 | 0.00 | -1516.3 |
| null |  | 2 | 3050.5 | 62.4 | 0.00 | -1523.3 |
| diel | hour | 5 | 3053.2 | 65.0 | 0.00 | -1521.6 |

Birds shifted downslope during ambient extremes. In model selection, the seasonal model was better than models that contained both hour and season covariates (Table 3). According to this seasonal model, birds were ∼4 m lower in the DS than the WS (*β*_dry_ = -4.35, SE = 0.90, *t* = - 4.85, *P* < 0.001, r^2^_c_ = 0.56). As individuals ranged across, on average, 41.2 m of vertical space in a given season (Appendix 3), 4 m constitutes an ∼10% proportional shift in vertical space use. In the second model (interaction between hour and season), bird elevation varied significantly by time of day within season (Figure 5B). Using 07:00 in the WS as the baseline, birds were ∼4 m higher at 16:00 in the WS (*β*_16:00, wet_ = 3.77, SE = 1.84, *t* = 2.05, *P* = 0.04) and ∼7 m lower at 16:00 in the DS (*β*_16:00, dry_ = -6.54, SE = 2.57, *t* = -2.54, *P* = 0.01, r^2^_c_ = 0.56). For the single bird tagged with a VHF radio tag, we collected 68 locations that were concentrated in the DS afternoon, but we saw the bird only once (1%) and triangulation suggested it often hid in a large, streamside log.

**Table 3.** Model selection for bird elevation, including the number of model parameters (K), AIC_c_ weight (*w*), and log likelihood (LL).

| model | covariates | K | AIC<sub>c</sub> | ΔAIC<sub>c</sub> | $w$ | LL |
| --- | --- | --- | --- | --- | --- | --- |
| season | season | 4 | 4965.3 | 0 | 0.87 | -2478.6 |
| interaction | hour*season | 10 | 4970.1 | 4.8 | 0.08 | -2474.9 |
| diel and season | hour + season | 7 | 4971.0 | 5.7 | 0.05 | -2478.4 |
| null |  | 3 | 4986.3 | 21.0 | 0.00 | -2490.1 |
| diel | hour | 6 | 4990.6 | 25.3 | 0.00 | -2489.3 |

Biologgers showed that birds altered their behavior by season (Figure 6). Light intensity varied considerably through time in both bird and ambient datasets—model r^2^ were relatively low, especially in the WS. Birds experienced a much darker environments than ambient biologgers indicated, with light exposure in the DS lower, even though light environment in the DS was far brighter. Temperature recorded by both bird and ambient loggers was higher in the DS, but the seasonal difference in peak temperature was about 50% lower for birds. Body temperatures of birds were high (41.7 ± 0.7°C, range 40.3 – 43.0°C, n = 36), with tags showing birds were exposed to higher ambient temperatures in the DS—tags recorded peaks at ∼36.3°C in the DS and ∼35.5°C in the WS. The timing of birds’ daily temperature peaks was similar in both seasons (14:16 and 14:21 in the DS and the WS, respectively). However, in contrast to ambient conditions, the variation in timing was usually smaller and similar across seasons for birds (Table 1).

**Figure 6.**
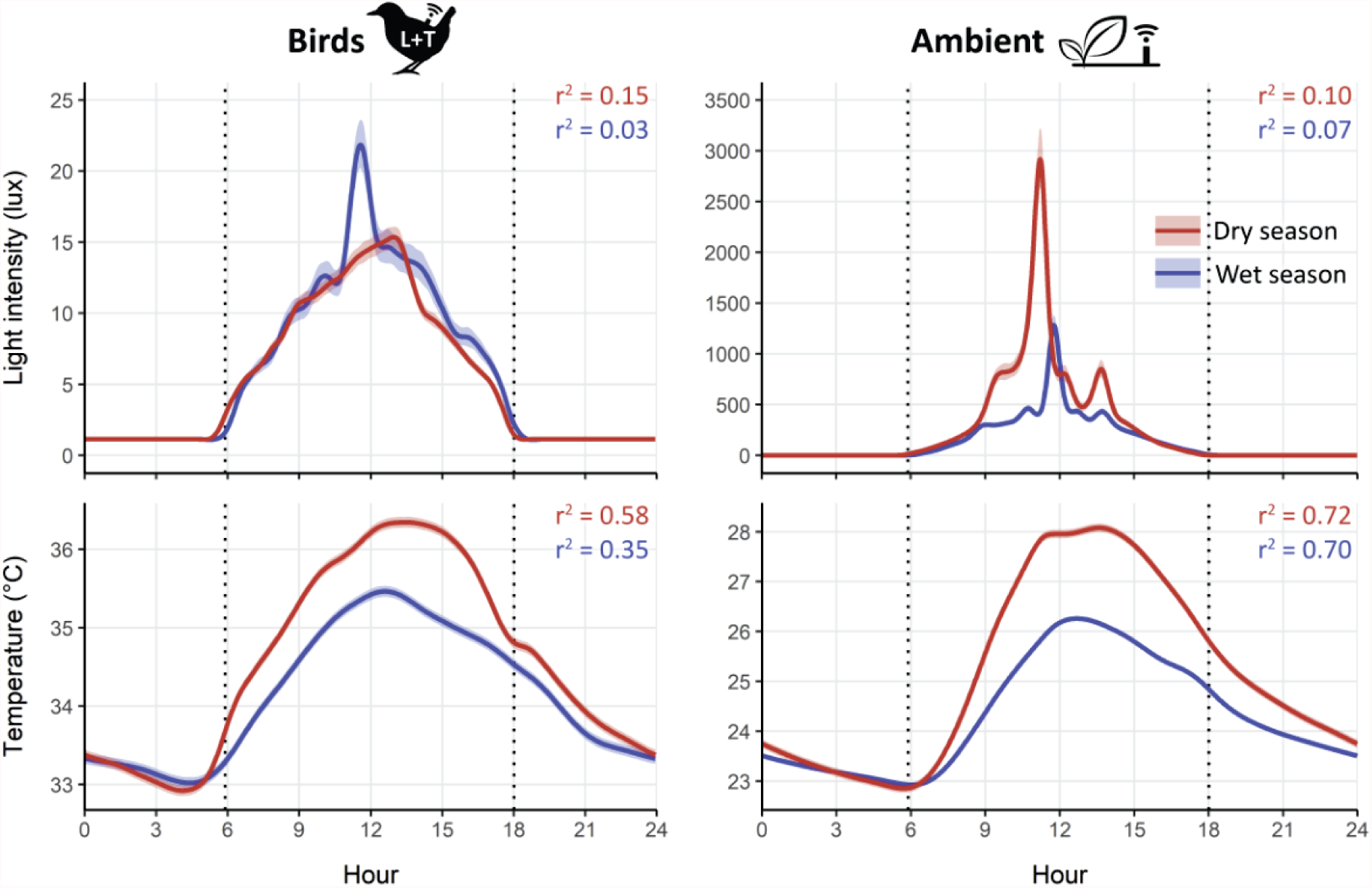
Light and temperature environment by day and season for birds and ambient conditions. Lines and 95% CIs are fits from generalized additive models with birds (*n* = 4) and ambient loggers (*n* = 4) as random effects. Note that y-axis on the ambient light plot is 2 orders of magnitude higher.

## DISCUSSION

We analyzed several high-resolution datasets for birds and their environment within Amazonian rainforest. We found that *F. analis*—a member of the sensitive terrestrial insectivore guild—attempted to sidestep extremes in ambient conditions within primary *terra firme* forest. Valleys and shelters offered microclimate refugia, and birds sought these during the afternoons of the dry season—the hottest and driest period of the year. Importantly, over the last four decades, climate change has made the dry season more extreme: we found that the modern dry season at the BDFFP is ∼21% drier and ∼1.3°C hotter than in the early 1980s—and these trends are only intensifying. Even if birds were not thermally stressed per se, periodic reductions in habitat and mobility constrain foraging opportunities; warming climate thus shortens the interval over which birds can meet their energetic needs (Bennett et al., 1984; Chappell & Bartholomew, 1981). Our study was motivated by alarming declines of terrestrial insectivores at both Amazonian sites with long-term data on primary forest avifauna, including the BDFFP (Blake & Loiselle, 2015; Stouffer et al., 2020). The ultimate goal of this study was to evaluate whether climate change is a possible mechanism. We found that climate is increasingly shifting towards conditions birds attempt to avoid, and despite evidence that birds actively influenced their thermal exposure, they managed to do so by only ∼50%. Our results are thus consistent with the hypothesis that climate change lowers habitat quality for terrestrial insectivores.

In lowland tropical rainforest, terrestrial insectivores inhabit the coolest and most stable of a warm environment, with a limited palette of options to behaviorally thermoregulate during hot and dry periods (Janzen, 1967). For these birds, we hypothesized that valleys and shelter may function as refugia and predicted that these would be used preferentially during ambient extremes. We then identified these periods empirically at two scales—daily and seasonal—and compared bird behavior at opposite ends of this ambient gradient. The top layer of soil, upon which terrestrial insectivores live and forage, experienced the highest temperatures between 14:00 and 15:00 in the DS (Table 1), but temperatures were lower in valleys (Figure 4). As expected, soil water content was related to rainfall—soils were drier in the DS and substantially drier at upper elevations than in valleys (Figure 4). Accordingly, we found that *F. analis* shifted its average elevation downslope during the DS—by up to 10 m (Figure 5B). Our proxy for shelter use (inverse probability of obtaining a GPS location) also revealed strong daily patterns within season (Figure 5A). Compared to a WS morning baseline, these results suggest that shelter use was 12x higher at 13:00 and 3x higher at 16:00 in the DS. Data from biologgers supported these results—although ambient light conditions were far brighter in the DS, birds at that time occupied darker areas (Figure 6). Still, birds experienced temperatures ∼1°C higher in the DS than the WS, though the ambient increase in DS temperature was ∼2°C.

A similar study from Panama did not find that nine understory insectivores selected buffered microclimates, or that microclimate varied appreciably over time (Pollock et al., 2015). However, that study was conducted in older secondary forest and utilized observers following radio-tagged individuals. We benefited from novel technology that let us sample on-bird conditions without observer disturbance and monitor the same individuals across seasons within primary forest. At our study site, we found stream temperature—within valleys in the DS—to be relatively constant and cool (24.6°C), which was 17.1°C lower than the average body temperature of *Formicarius analis* (41.7°C). Thus, even if only through bathing (Jullien & Thiollay, 1998), valley bottoms offer opportunities for rapid cooling.

Our data indicated that thermal and hydrological dynamics behaved as expected at the BDFFP, generating microclimate refugia at the valley bottoms. For instance, katabatic flow of colder air to low-lying areas (Pypker et al., 2007) lowers temperature in valleys, which tend to be already shaded. Hydrology within Amazonian micro-catchments is dominated by baseflow—rain infiltrates to groundwater and is slowly released into streams, which consequently have a relatively steady flow throughout the year (Tomasella et al., 2008), buffering temperature extremes (Davis et al., 2019; Elsenbeer, 2001; Fridley, 2009). More water within valleys may also allow cooling by enabling higher evapotranspiration when forest may be water-limited in drier areas and periods (Aleixo et al., 2019; Berg & Sheffield, 2019). Thus, in the DS, the demarcation between the dry upper slopes and wet lower slopes is not gradual (Figure 4). This threshold means that even a relatively small downslope shift can abruptly result in wet and cool conditions. In our data, the second DS was the only exception to this pattern—valleys showed higher maxima and range values, but we attributed this to treefalls that had occurred at two of the three low-elevation sites, likely leading to higher solar input. Otherwise, temperature average, minimum, maximum, and range were all higher in the DS and amplified on plateaus. Notably, the effect of elevation on temperature (but not water content) was erased in the WS. This is likely because cloud cover (Graham et al., 2003), air humidity (Aleixo et al., 2019), and environmental water content are all higher during the WS, mediating temperature fluctuations across the landscape. Passing clouds and precipitation possibly led to the inconsistent times of daily temperature peaks in the WS (Table 1).

Compared with downslope shifts, shelter use appears to be an additional but separate form of behavioral thermoregulation. Low rates of successful GPS fixes implied that birds sought cover in the afternoons of the DS—tags were nearly unable to acquire signal at 13:00 (3% success) and 16:00 (9% success), whereas the highest success rates occurred at 13:00 and 16:00 in the wet season (30 and 36%; Figure 5A). Elevation at bird locations suggested that birds shifted downslope between 13:00 and 16:00 in the DS, where GPS signal could be harder to acquire, but birds were not significantly lower when fix probability was lowest at 13:00. Our calibration data demonstrate that elevation alone cannot account for fix probability; both calibration tags obtained much higher success rates than bird tags, with the lower device attaining a slightly higher success rate than the upper device (69% vs 66%). This suggests that physical cover, in addition to valleys possibly blocking signal, drove the reductions in GPS fix rates. This notion was corroborated by our manually tracked bird—although observations concentrated in the afternoon, we only saw the bird once (1%) and triangulation suggested it often hid in a large, streamside log. Our results are thus consistent with the hypothesis that, during periods of ambient extremes, birds sought physical cover where ambient conditions were buffered (Scheffers et al., 2014).

We believe that our results are most compatible with birds responding directly to ambient extremes, but we cannot rule out indirect effects. Biomass of arthropod prey in the leaf litter drops with soil moisture (Jirinec et al., 2016; Levings & Windsor, 1984), including seasonal reductions in the DS (McKinnon et al., 2015; Pearson & Derr, 1986; Willis, 1976), and vertical movements that track moisture within the leaf litter (Usher, 1970). Mestre et al. (2010) quantified prey in regurgitated samples of *Formicarius analis* at the BDFFP, predominantly finding ants (Formicidae; ∼55%). In Panama, ant activity dropped by 25% in the DS and was >200% higher in ravines than exposed plateaus (Kaspari & Weiser, 2000). Capture rates of terrestrial insectivores as a guild—and *Formicarius analis* in particular—correlated with litter arthropod abundance, suggesting that these birds track resource availability within their home ranges (Karr & Brawn, 1990). Another terrestrial insectivore—the Song Wren (*Cyphorhinus phaeocephalus*)—was in the poorest body condition at the dry end of the rainfall gradient in Panama (Busch et al., 2011). Likewise, Spotted Antbirds (*Hylophylax naevioides*)—which, like *F. analis*, breed during the WS—were found to be ∼8-15% lighter during the late DS (Wikelski et al., 2000). Regardless of whether we detected a direct or indirect response to ambient extremes, the conclusion is likely the same—hot and dry periods are undesirable for terrestrial insectivores.

Future studies can augment this research in several ways. Although the variation in GPS fix success allowed us to estimate shelter use, the challenging environment for GPS tags resulted in a loss of ∼75% of locations with elevation data. This introduced two issues. First, the markedly smaller size of the elevation dataset may have reduced our ability to resolve the effect of season and daytime as AIC_c_ substantially penalized the interaction model due to its complexity (Table 2). Second, areas and times where tags received relatively few fixes may have been underrepresented—an issue raised in previous studies on habitat selection (D’Eon, 2003). However, GPS tags recorded substantially higher number of fixes in the WS when GPS signal should have been hindered by cloud cover (Graham et al., 2003; Figure 6). To obtain these results, birds must have moved higher up and to more open locations during the WS. Researchers could avoid the above issues with direct observations of radio-tagged birds, though that raises concerns about logistics and observer effects on birds—both potentially insurmountable. Overall, our conclusions stand on relatively high-resolution and diverse datasets, but we only considered a single species and two seasonal cycles. Nevertheless, the ultimate cause of biological seasonality is climate over evolutionary time. We propose two studies that could provide further evidence for negative influence of climate change: 1) testing if less sensitive ground-foraging species do not respond to ambient extremes, and 2) a decadal study showing that annual elevational shifts and shelter use are a function of the seasonal severity for a given year (e.g., more extreme response during hot and dry El Niño years).

## CONCLUSIONS

This study highlights that heterogeneity in the ‘stable’ lowland rainforest exists in three domains—daily, seasonal, and topographical. *Formicarius analis*, a member of the terrestrial insectivore guild, which should be among the most buffered from periodic fluctuations in temperature and water availability, nonetheless seeks shelter and moves to areas where conditions are milder during ambient extremes. Regardless of whether our study revealed that these birds behaviorally thermoregulate or that they instead track a shifting food base (or both), climate change exacerbates the very conditions that lead to such behavior and thus likely shrinks suitable habitat for this species at the BDFFP. *Formicarius analis* and other terrestrial insectivores may thus be especially vulnerable to climate change. In fact, the impetus for this study was the substantial decline of terrestrial insectivores reported recently within primary forests at the BDFFP (Stouffer et al., 2020). Previous decades of tracing avian responses to forest fragmentation raise the possibility that hotter and drier conditions may be at least partially responsible for disappearances in fragments. Our results are consistent with the predictions of this microclimate hypothesis for birds in continuous primary forest. Furthermore, we underscore that climate change will increasingly produce such conditions in lowland Amazonia, which mostly lacks topographical variation and associated refugia. If these sensitive specialists act as a barometer within the vast and biodiverse forests of Amazonia, their behavior raises cause for concern.

## ACKNOWLEDGEMENTS

We thank Bruna Amaral, Flamarion Assunção, and Jairo Lopes for tireless assistance in the field. Erik Johnson, Richard Keim, Michael Kaller, Michael Dance, and Stephen Midway provided constructive comments in analysis and interpretation of data. Ryan Burner compiled ERA5-Land data for climate analyses. Field logistics were facilitated by the BDFFP support personnel, especially Rosely Hipólito, Manoela Borges, José Luís Camargo, Mario Cohn-Haft, and Ary Ferreira. Funding for this research was provided by the US National Science Foundation (LTREB 0545491 and 1257340), the National Institute of Food and Agriculture, US Department of Agriculture, McIntire Stennis projects no. 94098 and no. 94327, Smithsonian Tropical Research Institute Short-Term Fellowship, Neotropical Bird Club Conservation Fund, Lewis and Clark Fund for Exploration and Field Research from the American Philosophical Society, and research grants from the American Ornithological Society, Animal Behavior Society, and the Wilson Ornithological Society. This is publication no. XXX of the BDFFP Technical Series and no. XX of the Amazonian Ornithology Technical Series of the INPA Collections Program. The manuscript was approved by the Director of the Louisiana State University Agricultural Center as manuscript number XXXX-XXX-XXXX.

## APPENDIX 1

**Appendix 1.**
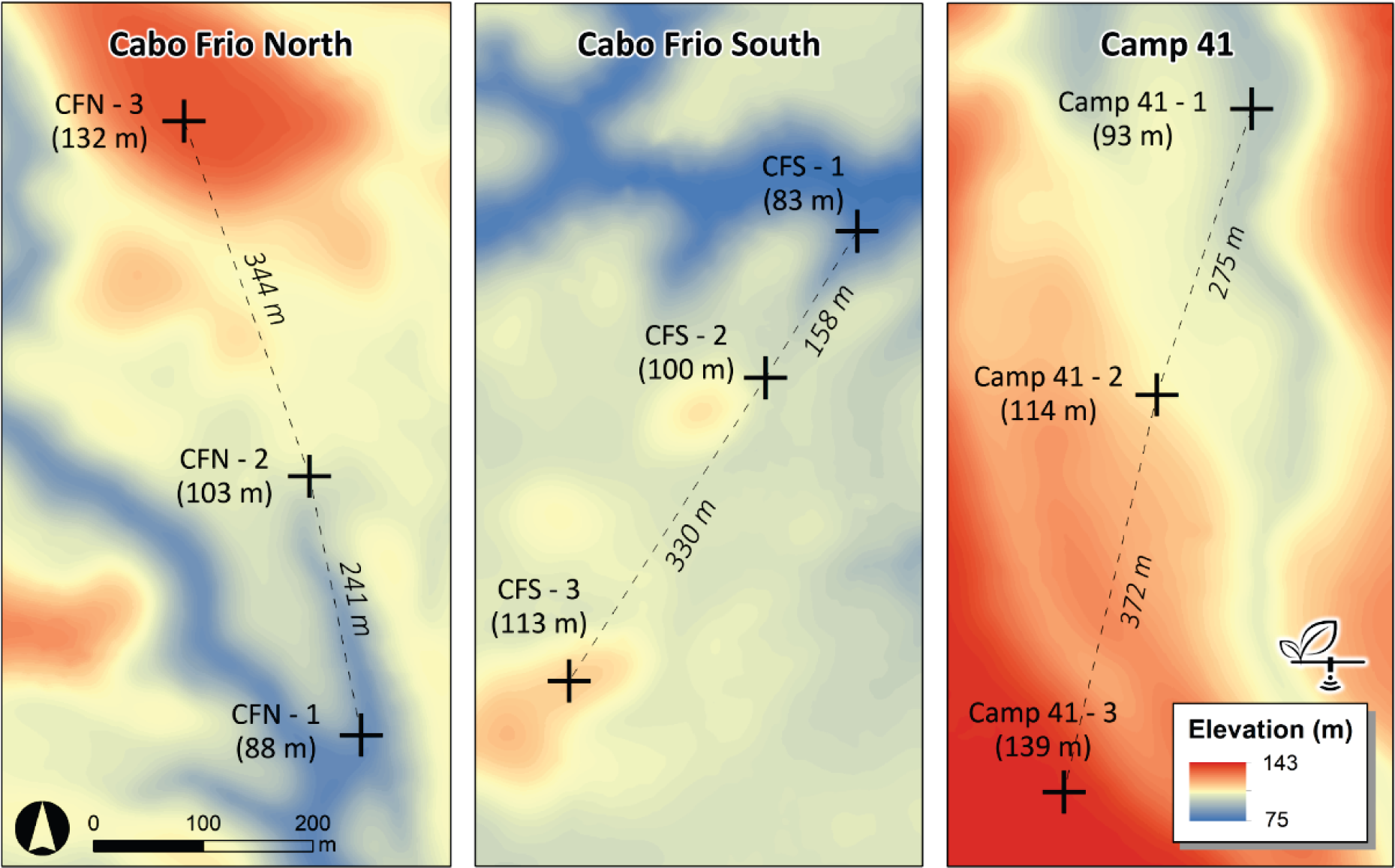
Microclimate datalogging transects within Amazonian micro-catchments. We placed nine stations (black crosses) along three transects, each logging soil temperature and moisture data every 10 min from June 2017 to September 2019.

## APPENDIX 2

**Appendix 2.**
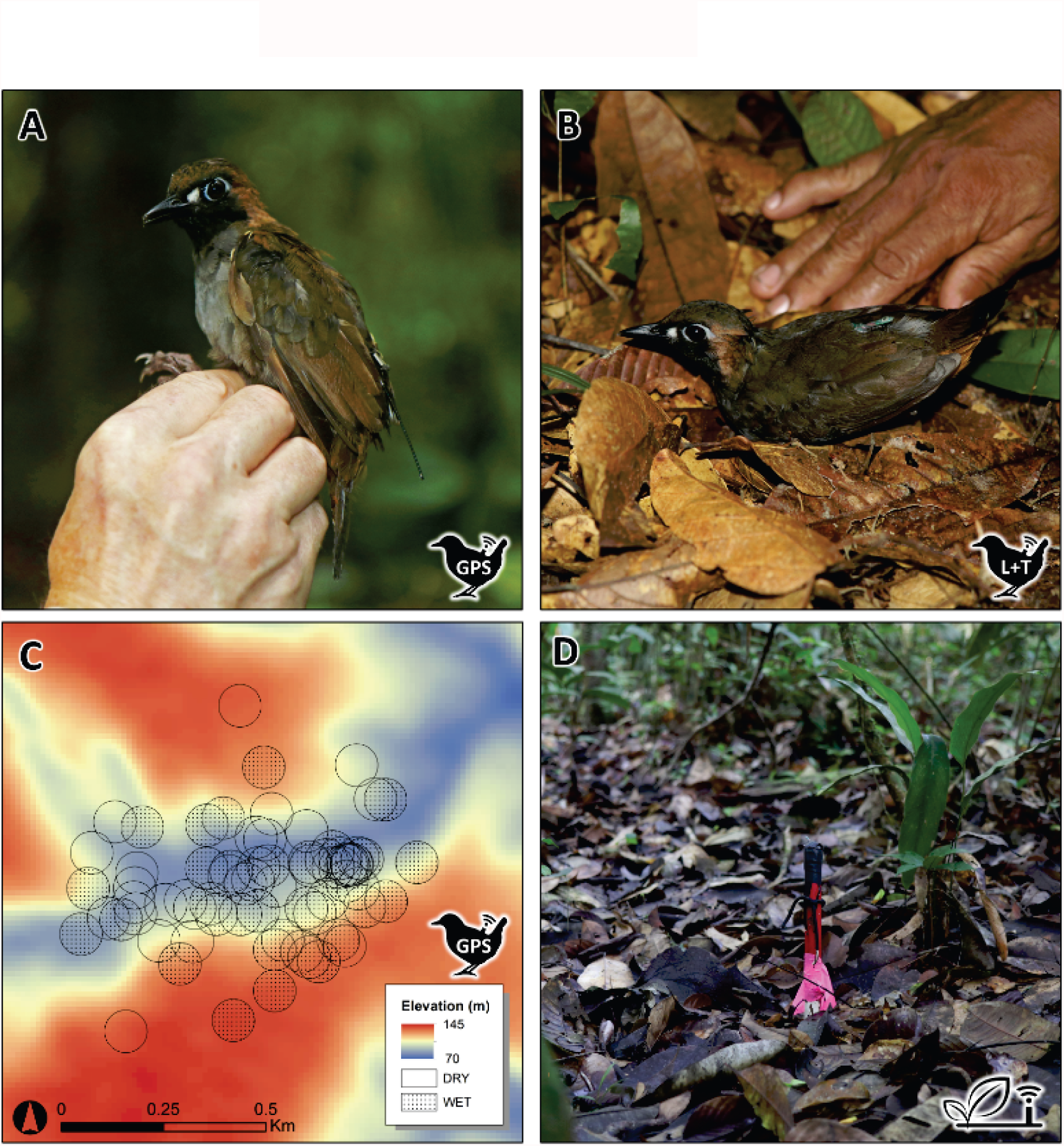
Tracking *Formicarius analis*. Birds were outfitted with either a GPS tag that recorded locations (A), or a biologger that recorded light intensity and temperature (B). Panel (C) shows locations for one bird categorized by season (dry, wet), where we averaged elevation values within 52 m buffers to account for location error estimated by calibration tags. Calibration tags (both GPS and biologgers) were placed atop short stakes within the forest understory to sample ambient conditions (D).

## APPENDIX 3

**Appendix 3.**
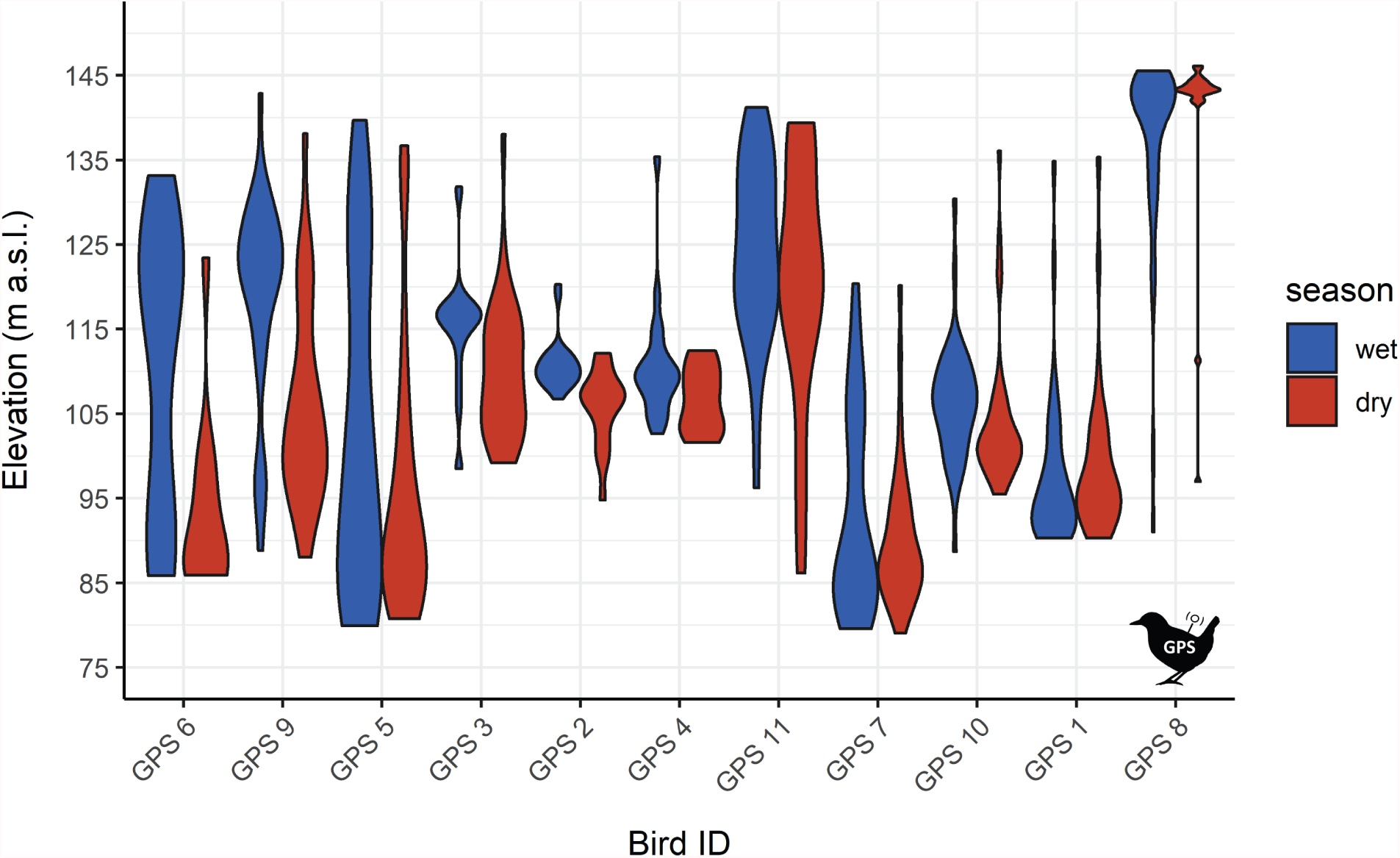
Violin plots of individual bird elevations by season. Birds are ordered by the mean elevation reduction in the dry season.

